## Supplementary data for "Structure of the turnover-ready state of an ancestral respiratory complex I"

**Extended Data**

**Extended Data Table 1. Cryo-EM data collection, refinement, and validation statistics for *Pd*-CI-DDM.**

| **Data collection and processing** | ***Pd*-CI-DDM** | |
| --- | --- | --- |
| Voltage (kV) | 300 | |
| Nominal magnification | 130,000× | |
| Electron exposure (*e^–^* Å^-2^) | 48 | |
| Defocus range (μm) | -1.5 to -2.9 | |
| Calibrated pixel size (Å) | 1.05 | |
| Number of frames | 25 | |
| Energy filter slit width (eV) | 20 | |
| Symmetry imposed | C1 | |
| Number of micrographs | 2,278 | |
| Initial particle images (no.) | 128,270 | |
| Final particle images (no.) | 51,308 | |
| **Final Global Classification** | **Class 1** (EMDB: 19975) | **Class 2** (EMDB: 19976) |
| Final particle images (no.) | 19,736 | 31,572 |
| Map sharpening *B* factor (Å^2^) | -76.26 | -95.39 |
| Map resolution (Å) (FSC = 0.143) | 4.5 | 4.2 |
| Map resolution range (Å) | 3.75 – 12.65 | 3.99 – 12.55 |

**Extended Data Table 2. Summary of the model for *Pd*-CI-ND** (PDB-8QBY).

| *Pd*-CI subunit | Human CI  subunit | Chain identity | Total residues | Modelled  residues (%) | Q-score | Cofactors & modification |
| --- | --- | --- | --- | --- | --- | --- |
| Nqo1 | NDUFV1 | F | 431 | 1-421 (97.7) | 0.83 | FMN, 4Fe4S |
| Nqo2 | NDUFV2 | E | 239 | 1-236 (98.7) | 0.83 | 2Fe2S |
| Nqo3 | NDUFS1 | G | 674 | 2-667 (98.8) | 0.84 | 2Fe2S, 2 x 4Fe4S, Na^+^ |
| Nqo4 | NDUFS2 | D | 412 | 3-412 (99.5) | 0.87 | Dimethyl-Arg65, Ca^2+^ |
| Nqo5 | NDUFS3 | C | 208 | 6-196 (91.8) | 0.86 |  |
| Nqo6 | NDUFS7 | B | 175 | 28-175 (84.6) | 0.87 | 4Fe4S |
| Nqo9 | NDUFS8 | I | 163 | 3-163 (98.8) | 0.87 | 2 x 4Fe4S |
| Nqo8 | ND1 | H | 345 | 1-342 (99.1) | 0.86 |  |
| Nqo14 | ND2 | N | 499 | 1-479 (96.0) | 0.86 | Ca^2+^ |
| Nqo7 | ND3 | A | 121 | 1-121 (100) | 0.86 | N-formyl |
| Nqo13 | ND4 | M | 513 | 1-503 (98.1) | 0.83, 0.83^2^ | Ca^2+^ |
| Nqo11 | ND4L | K | 101 | 1-101 (100) | 0.86 |  |
| Nqo12 | ND5 | L | 703 | 1-506, 551-703 (93.7) | 0.78, 0.82^2^ |  |
| Nqo10 | ND6 | J | 200^1^ | 1-82, 88-200 (97.5) | 0.84 | N-terminus 2 x Met |
| *Pd*NUYM | NDUFS4 | Q | 103 | 1-103 (100) | 0.84 |  |
| *Pd*NUMM | NDUFS6 | R | 62 | 3-61 (95.2) | 0.83 | Zn^2+^, Ca^2+^ |
| *Pd*N7BM | NDUFA12 | q | 124 | 1-124 (100) | 0.84 |  |
| PIMT | - | t | 217 | 2-217 (99.5) | 0.83 |  |

^1^The total number of residues for ND6 is 200, taking into account the two Met residues observed at the N-terminus

^2^Q-score determined for the focused map of the membrane ND4-ND5 subdomain

**Extended Data Table 3. Sequence and structural comparison of *Pd*-CI to other CI species.**

| Subunit | Relative to *B. taurus* complex I subunits | | |  | |
| --- | --- | --- | --- | --- | --- |
|  | Sequence identity (%) | Length (%)  (N of residues) | RMSD^1^ values (Å) | | |
|  |  |  | *B. taurus* | *Y. lipolytica* | *E. coli* |
| NDUFV1 (Nqo1) | 64.0 | 92.9 (-33) | 0.60 | 0.89 | 2.34 |
| NDUFV2 (Nqo2) | 40.2 | 96.0 (-10) | 6.66 | 9.00 | 1.35 |
| NDUFS1 (Nqo3) | 47.5 | 92.7 (-53) | 3.14 | 2.25 | 11.68 |
| NDUFS1 N/C-domains | 61.9/40.5 | 92.7 (-18/-35) | 1.07/3.89 | 1.11/2.55 | 3.49/13.97 |
| NDUFS2 (Nqo4) | 59.0 | 89.0 (-51) | 5.17 | 5.69 | 2.72 |
| NDUFS3 (Nqo5) | 46.9 | 78.2 (-58) | 1.97 | 1.51 | 19.89 |
| NDUFS7 (Nqo6) | 68.0 | 81.0 (-41) | 1.13 | 1.43 | 1.41 |
| NDUFS8 (Nqo9) | 73.0 | 76.9 (-49) | 0.80 | 1.07 | 3.20 |
| ND1 (Nqo8) | 39.1 | 108.5 (+27) | 2.51 | 3.25 | 1.81 |
| ND2 (Nqo14) | 25.7 | 143.8 (+152) | 4.51 | 3.75 | 3.31 |
| ND3 (Nqo7) | 37.5 | 105.2 (+6) | 1.31 | 3.57 | 2.30 |
| ND4 (Nqo13) | 31.4 | 111.8 (+54) | 3.01 | 4.81 | 2.32 |
| ND4L (Nqo11) | 22.9 | 103.1 (+3) | 1.64 | 1.73 | 1.17 |
| ND5 (Nqo12) | 34.7 | 116.0 (+97) | 4.09 | 3.48 | 4.05 |
| ND6 (Nqo10) | 23.6 | 114.3 (+25) | 7.25 | 6.99 | 4.77 |
| NDUFS4 (dNUYM) | 36.0 | 58.9 (-72) | 2.51 | 3.03 | - |
| NDUFS6 (PdNUMM) | 32.7 | 50.0 (-62) | 2.21 | 4.27 | - |
| NDUFA12 (PdN7BM) | 37.0 | 85.5 (-21) | 9.56 | 8.94 | - |
| PIMT | - | - | - | - | - |

^1^The structures used for the RMSD comparisons were PDB-7QSL (2.8 Å-resolution bovine structure in the closed state), PDB-6YG4 (2.7 Å-resolution *Y. lipolytica* structure) and PDB-7Z7S (2.4 Å-resolution *E. coli* structure in the closed state). RMSD values were calculated using UCSF ChimeraX including all pairs of residues.

**Extended Data Table 4. Structural investigation of the inter-domain angle of complex I.**

|  | Species | Given name | PDB | Interdomain angle^1^ (°) | | |
| --- | --- | --- | --- | --- | --- | --- |
|  |  |  |  | 4Fe-ND1-ND5 | 4Fe-ND1-ND2 | N2-ND1-ND4L |
| Bacteria | *Paracoccus denitrificans* | Active/closed | this  work | 117.0 | 124.2 | 127.5 |
|  | *Mycobacterium smegmatis* | Active | 8E9G | 118.3 | 126.0 | 128.8 |
|  | *Escherichia*  *coli* | Closed | 7Z7S | 118.7 | 119.2 | 126.9 |
| Single cell eukaryote | *Tetrahymena thermophila* | Closed | 7TGH | 112.4 | 121.1 | 130.7 |
| Insect | *Drosophila melanogaster* | Active (closed) | 8B9Z | 113.1 | 120.5 | 127.6 |
| Mammal | *Bos*  *taurus* | Active-apo (closed) | 7QSL | 112.0 | 120.3 | 120.4 |
|  |  | Deactive-apo (open) | 7QSN | 113.5 | 121.2 | 125.3 |
|  | | | | 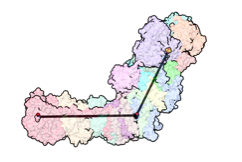 | 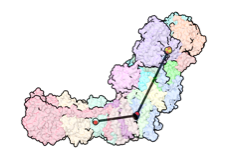 | 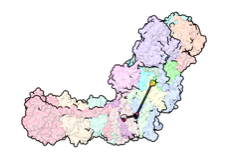 |

^1^The conserved reference points chosen to measure the inter-arm angles were the N2 [4Fe-4S] cluster in NDUFS7, the 4Fe all-Cys ligated [4Fe-4S] cluster in NDUFS1 and the Cα of *Pd*-ND1-L126, *Pd*-ND5-F349 and *Pd*-ND4L-E36. Angles were measured using UCSF ChimeraX. The structures used in this analysis were PDB-8E9G, PDB-7Z7S, PDB-7TGH, PDB-8B9Z, PDB-7QSL and PDB-7QSN.

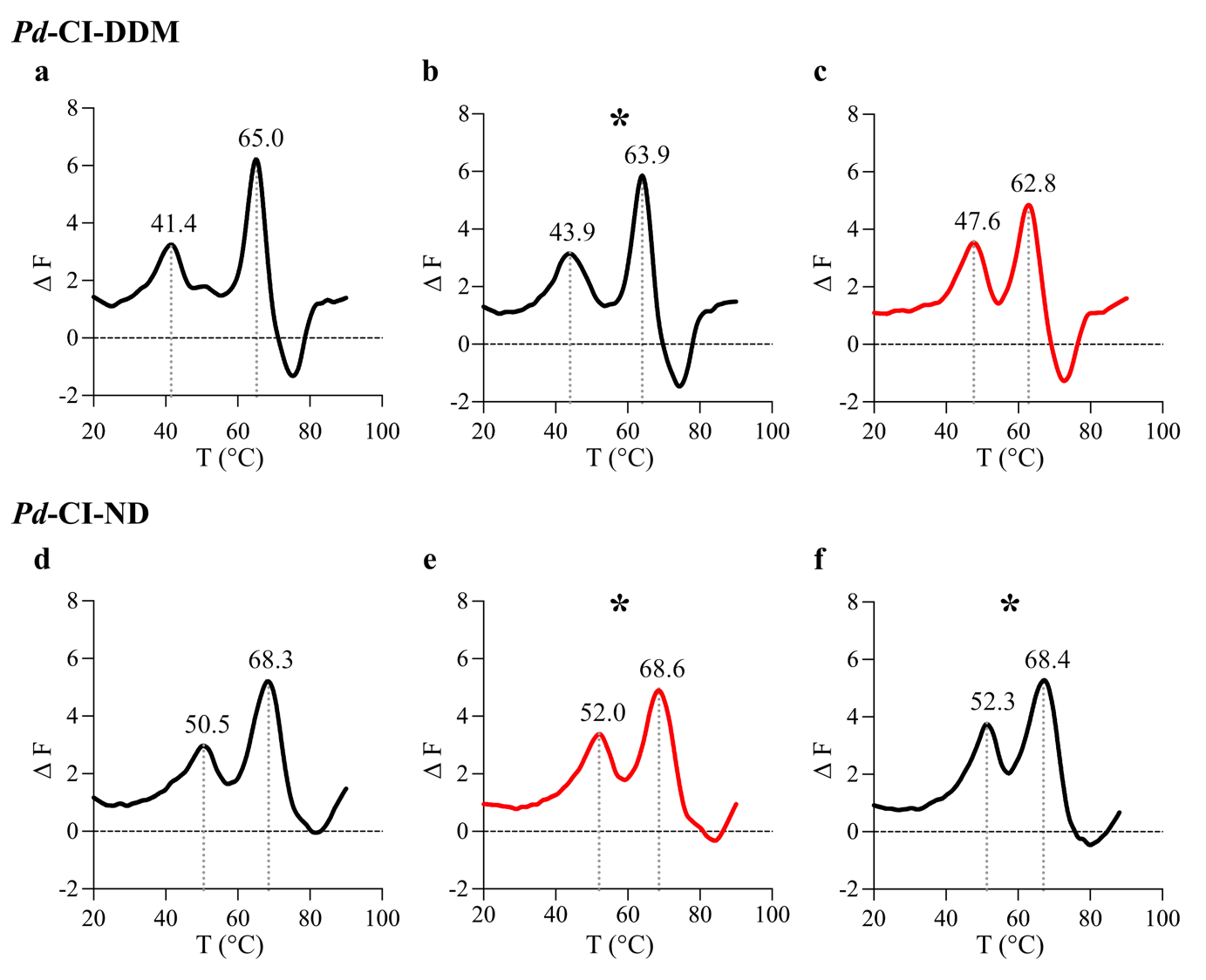

**Extended Data Figure 1. Comparison of the stability of *Pd*-CI-DDM and *Pd*-CI-ND preparations by nano-DSF.** The data are presented as the first derivative of the change in fluorescence as the temperature is increased with peak positions labelled. a) *Pd*-CI-DDM in the buffer used for size-exclusion chromatography (20 mM MES (pH 6.5 at 4 °C), 150 mM NaCl, 10 mM CaCl_2_, 10% (v/v) glycerol, 0.05% (w/v) DDM). b) *Pd*-CI-DDM in the buffer for cryo-EM grid preparation (the buffer from (a) but without glycerol). c) *Pd*-CI-DDM in 20 mM MES (pH 6.5 at 4 °C), 25 mM NaCl, 1 mM CaCl_2_ and 0.05% DDM. d) *Pd*-CI-ND in ND reconstitution buffer (20 mM MES (pH 6.5 at 4 °C) and 25 mM NaCl). e) *Pd*-CI-ND in the buffer for cryo-EM grid preparation (ND reconstitution buffer plus 1 mM CaCl_2_). f) *Pd*-CI-ND with FOM in the buffer for cryo-EM grid preparation (ND reconstitution buffer plus 1 mM CaCl_2_ and 0.01% FOM). Panels (c) and (e) (red) directly compare *Pd*-CI-DDM and *Pd*-CI-ND under matching conditions, and asterisks denote conditions that were used to make cryo-EM grids. Each trace was repeated three times, and the peak positions were consistent within 0.5 °C ranges.

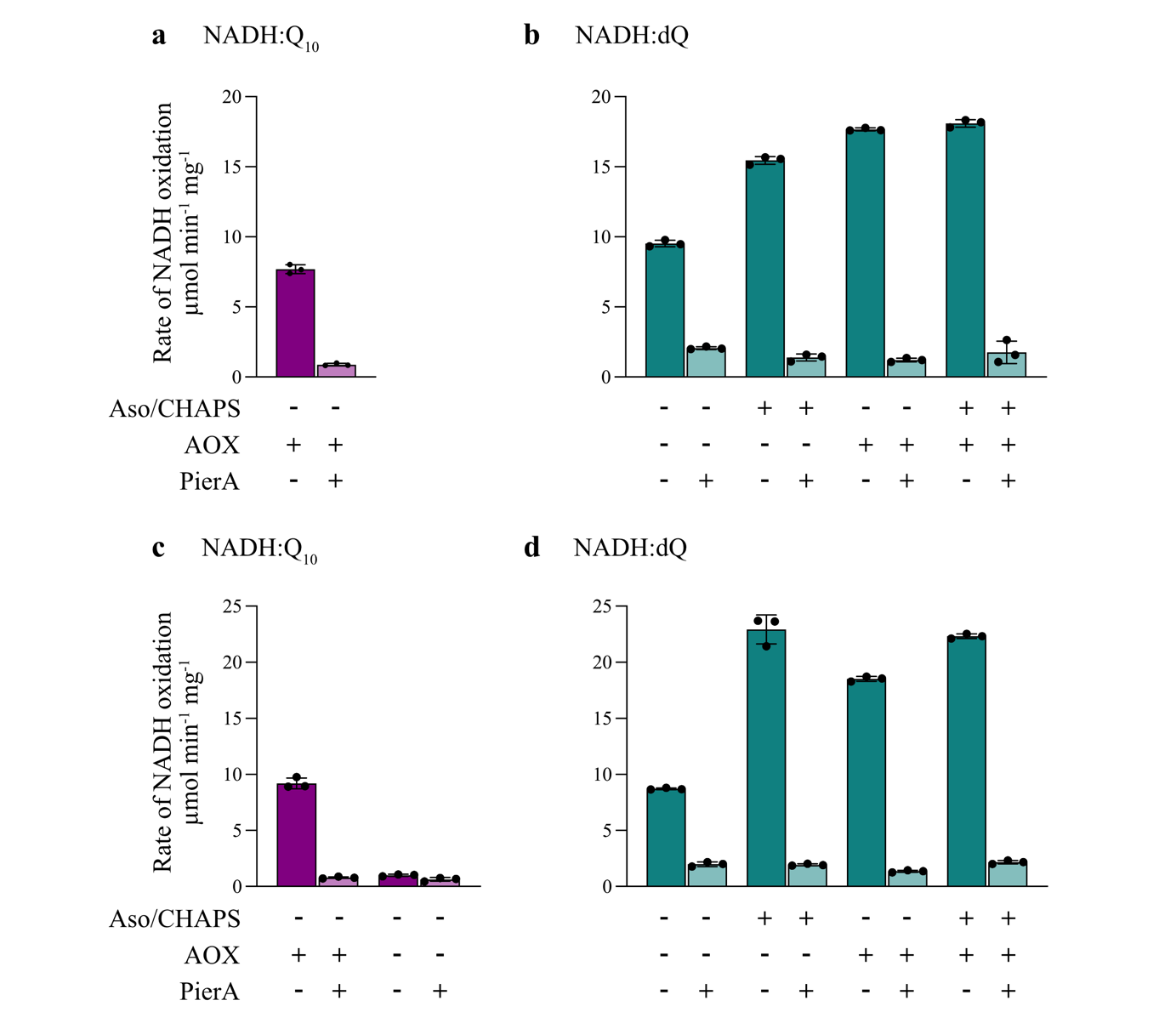

**Extended Data Figure 2. Catalytic activities of the two *Pd*-CI-ND preparations analyzed by cryo-EM.** a) Rates of NADH:ubiquinone-10 oxidoreduction using the ubiquinone-10 contained in the nanodiscs and recycled by AOX. b) Rate of NADH:dQ oxidoreduction by *Pd*-CI-ND (grids 1 and 2). Assays were carried out using 200 μM dQ as described in *Materials and Methods*, with the addition of asolectin and CHAPS to solubilise the nanodiscs, AOX to recycle ubiquinol, and piericidin A to inhibit catalysis as indicated. c) and d) Equivalent data for *Pd*-CI-ND (grid 3) frozen in the presence of FOM. All rates have been normalised for the complex I content determined using the NADH:APAD^+^ oxidoreduction assay. The activities of the two *Pd*-CI-DDM preparations before reconstitution were 22.0 ± 1.9 (grids 1 and 2) and 19.5 ± 0.2 µmol min^−1^ mg^−1^ (grid 3).

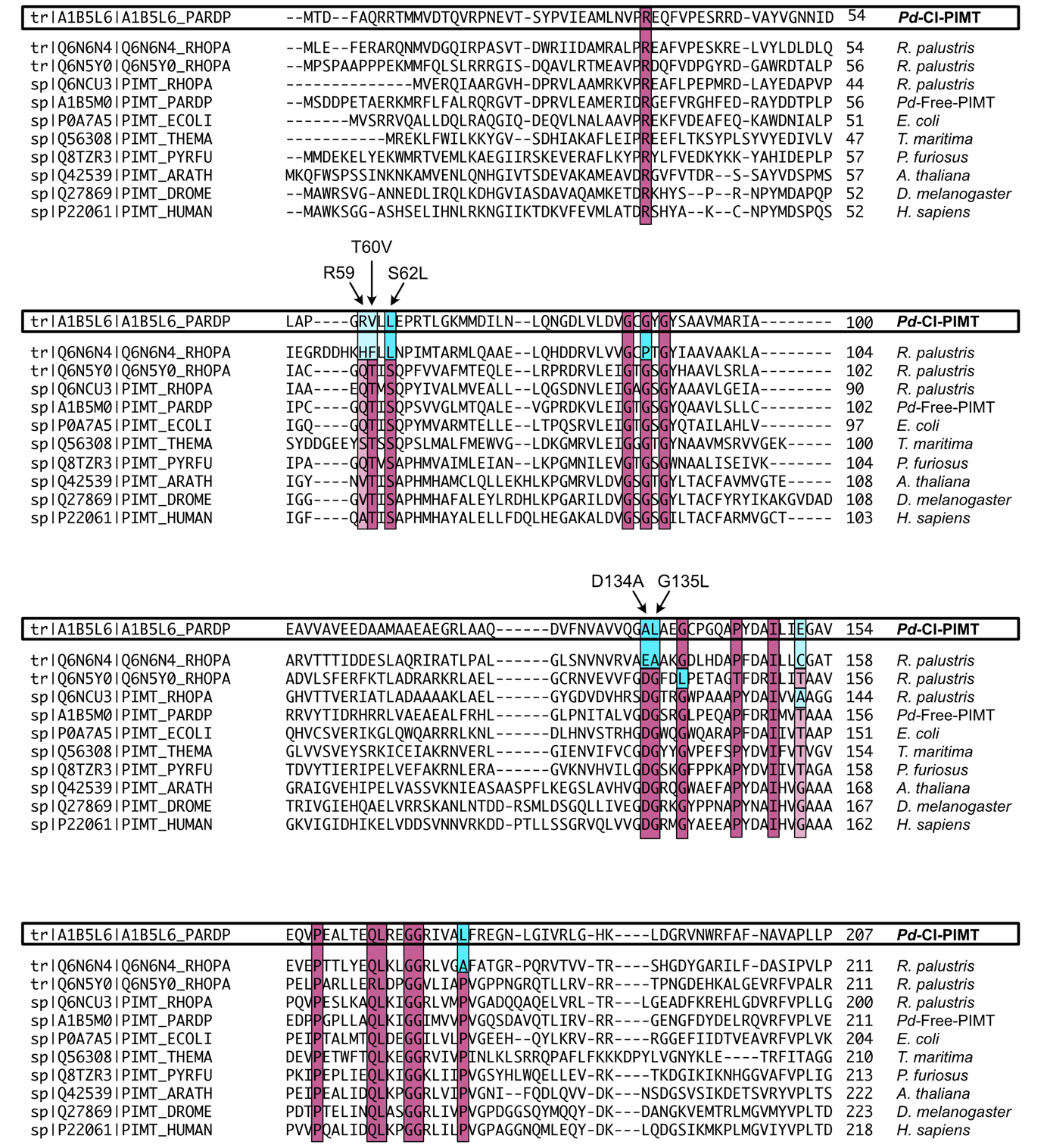

**Extended Data Figure 3. Sequence alignment of PIMT across different species.** Figure showing the sequence alignment of *Pd*-CI-PIMT (A1B5L6) against the free PIMT paralog from *Pd*-CI (A1B5M0) and orthologues from *Rhodopseudomonas palustris* (Q6N6N4, Q6N5Y0, Q6NCU3), *Escherichia coli* (P0A7A5)*, Thermotoga maritima* (Q56308)*, Pyrococcus furiosus* (Q8TZR3)*, Arabidopsis thaliana* (Q42539), *Drosophila melanogaster* (Q27869) *and Homo sapiens* (P22061). Universally conserved residues are highlighted in maroon while divergent mutations in *Pd*-CI-PIMT and *R. palustris* Q6N6N4 are shown in cyan. Lighter shades indicate small side-chain amino acids residues that are mutated to bulkier residues in *Pd*-CI-PIMT to fill the space occupied by the permanently bound SAM cofactor in functional PIMTs.

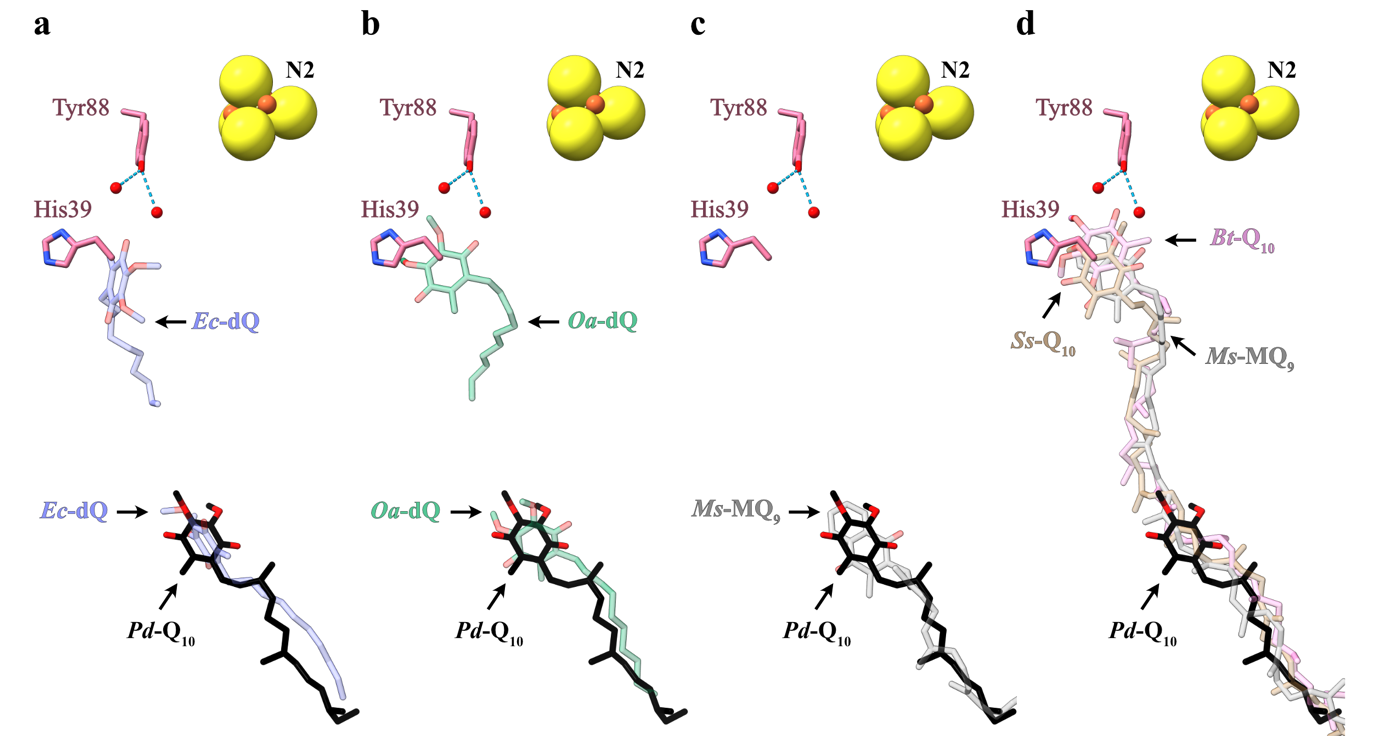

**Extended Data Figure 4. Comparison of closed structures with Q_10_, menaquinone-9 (MQ_9_) and short-chain decylubiquinone (dQ) bound to the Q_10_-bound structure of *Pd*-CI.** a) The dQ bound in *Ec-*CI (blue, PDB-7Z7S) overlayed with the partially bound Q_10_ in *Pd*-CI*.* b) Comparison of the fully and partially inserted dQ molecules in ovine CI (green, PDB-6ZKC) with the Q_10_ bound in *Pd*-CI. c) Partially bound MQ_9_ in *M. smegmatis* CI (grey, PDB-8E9G) overlayed with partially bound Q_10_ in *Pd*-CI. d) Fully inserted Q_10_ or MQ_9_ positions in bovine (pink, PDB-7QSK), porcine (wheat, PDB-7V2C) and *M. smegmatis* complex I showing the ~20 Å distance between the two Q head groups binding positions.

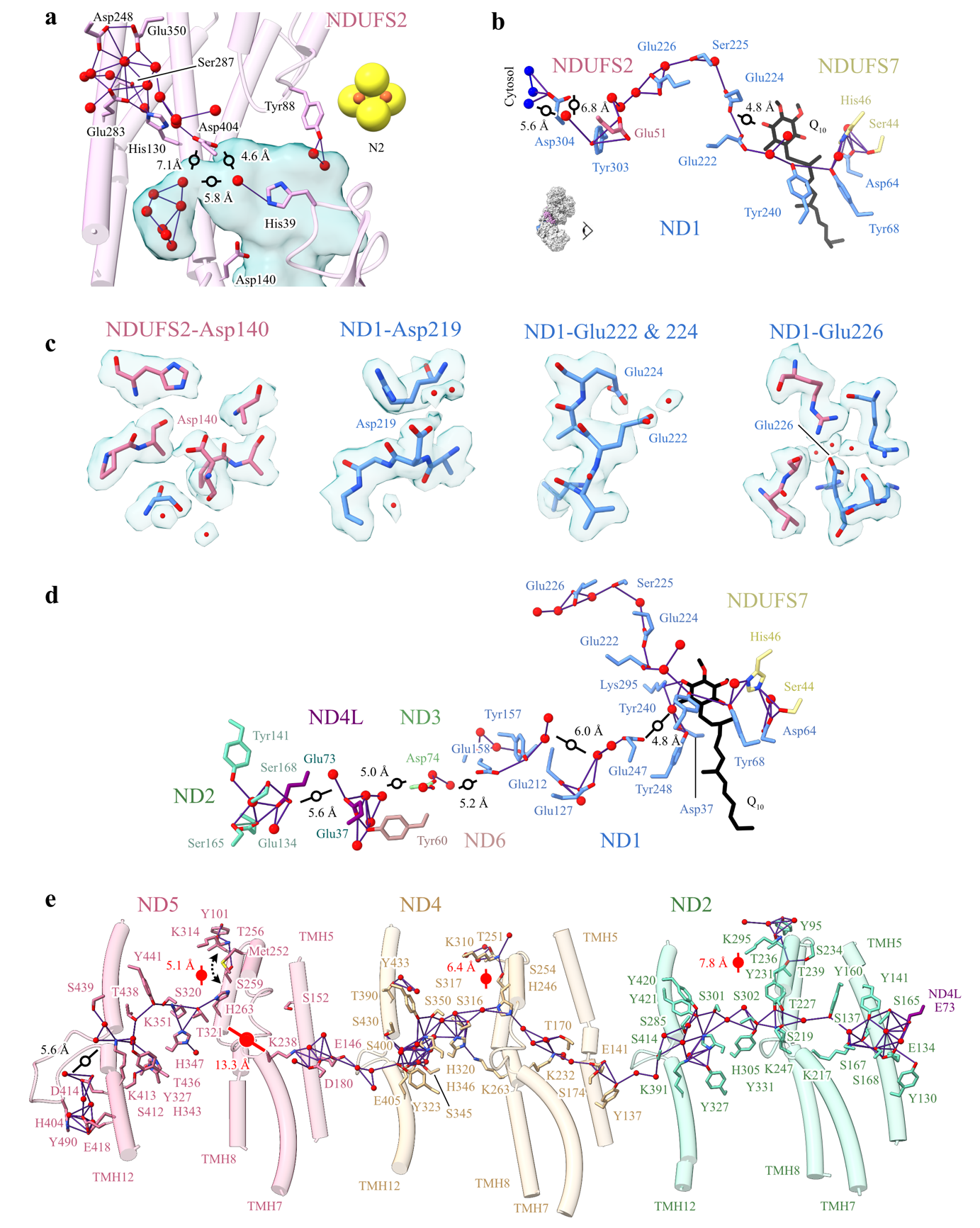

**Extended Data Figure 5. Grotthuss competent pathways in *P. denitrificans* complex I at the ubiquinone site, E-channel, and central axis.** a) A potential ubiquinone-10 protonation pathway leading to the ubiquinone-binding active site. b) A second potential ubiquinone-10 protonation pathway leading to the secondary Q-binding site. c) The locally sharpened cryo-EM densities of key acidic residues. Panels show the cryo-EM density outline at a 1.5 Å range from the displayed atoms as a semi-transparent surface displayed in UCSF ChimeraX at map threshold of 0.01. Each acidic residue is displayed with residues or water molecules within 3 Å distance. d) Grotthuss competent network of the lower section of the ubiquinone-10 binding site and the E-channel. e) Grotthuss competent networks in the ND2, ND4 and ND5 subunits in *Pd*-CI. The networks are shown with purple lines and the gaps are indicated by black lines (with distances displayed) and open circles highlighting protein unobstructed routes. Red lines with full circles indicate protein obstructed gaps.

**Supplementary Data**

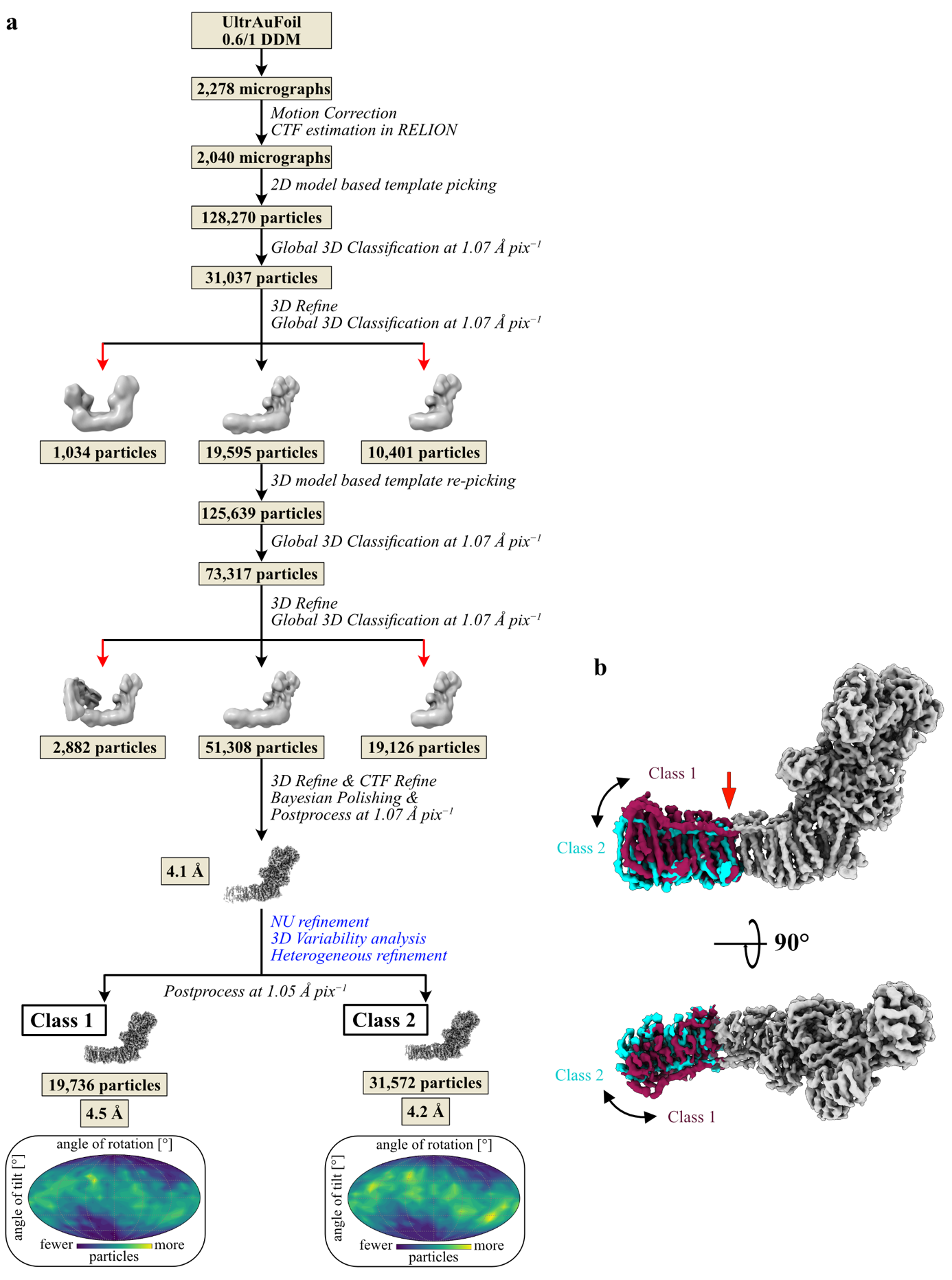

**Figure S1. Cryo-EM data processing pipeline for *Pd*-CI-DDM and comparison of classes 1 and 2.** a) Cryo-EM data processing pipeline, with the red arrows indicating particles that were excluded. The orientation distributions for each class are shown. b) Comparison of the density maps for classes 1 and 2 aligned on the hydrophilic domain showing the different global conformations of the membrane domain. The cleft that opens up in class 2 between subunits ND2 and ND4, allowing the ND4-5 distal subdomain to move relative to the rest of the enzyme, is marked with an arrow. Processing steps in blue were carried out in *CryoSPARC* *v3.3.2*.

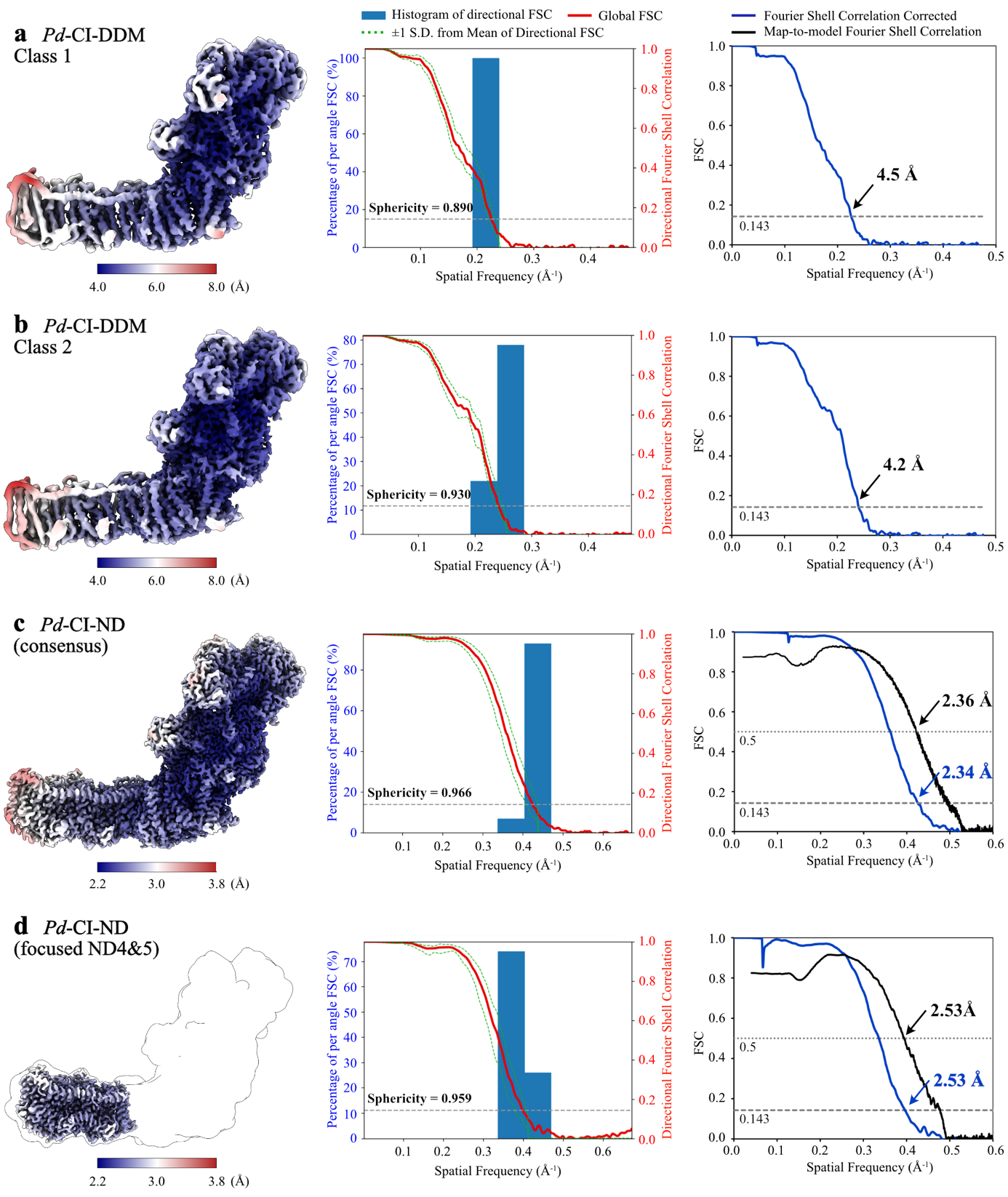

**Figure S2. Local resolution and FSC curves for all *Pd*-CI maps.** Local resolution plots calculated in RELION and visualised in UCSF ChimeraX (left) are shown with 3D FSC curves and histograms showing the distribution of resolution shells (middle) and half-map/map-to-model FSC curves (right) for a) *Pd*-CI-DDM class 1; b) *Pd*-CI-DDM class 2; c) *Pd*-CI-ND (consensus); and d) *Pd*-CI-ND focussed map for the distal membrane subdomain.

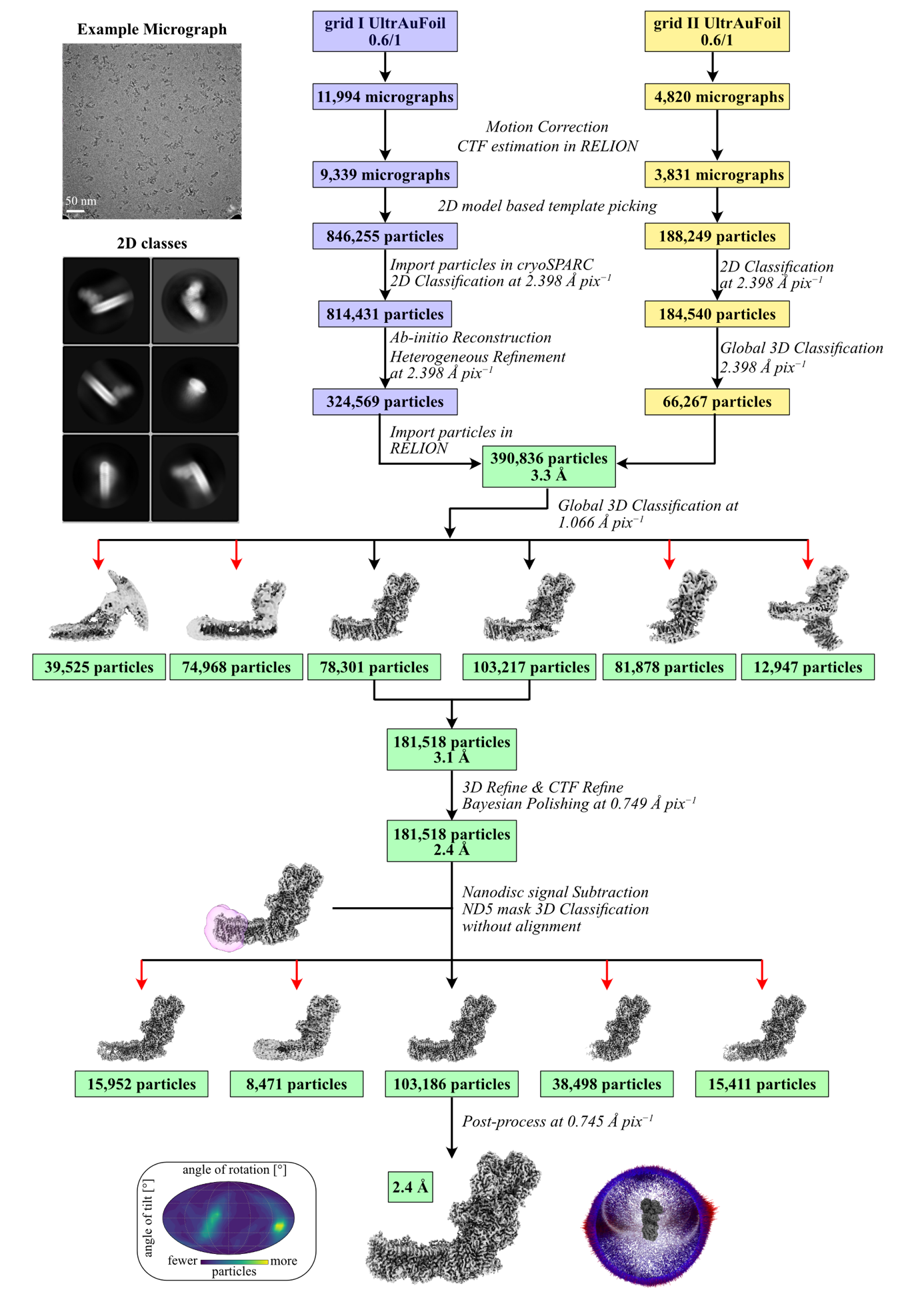

**Figure S3. Cryo-EM data processing pipeline for grids 1 and 2 for *Pd*-CI-ND.** The red arrows indicate particles that were excluded, and the orientation distributions are shown.

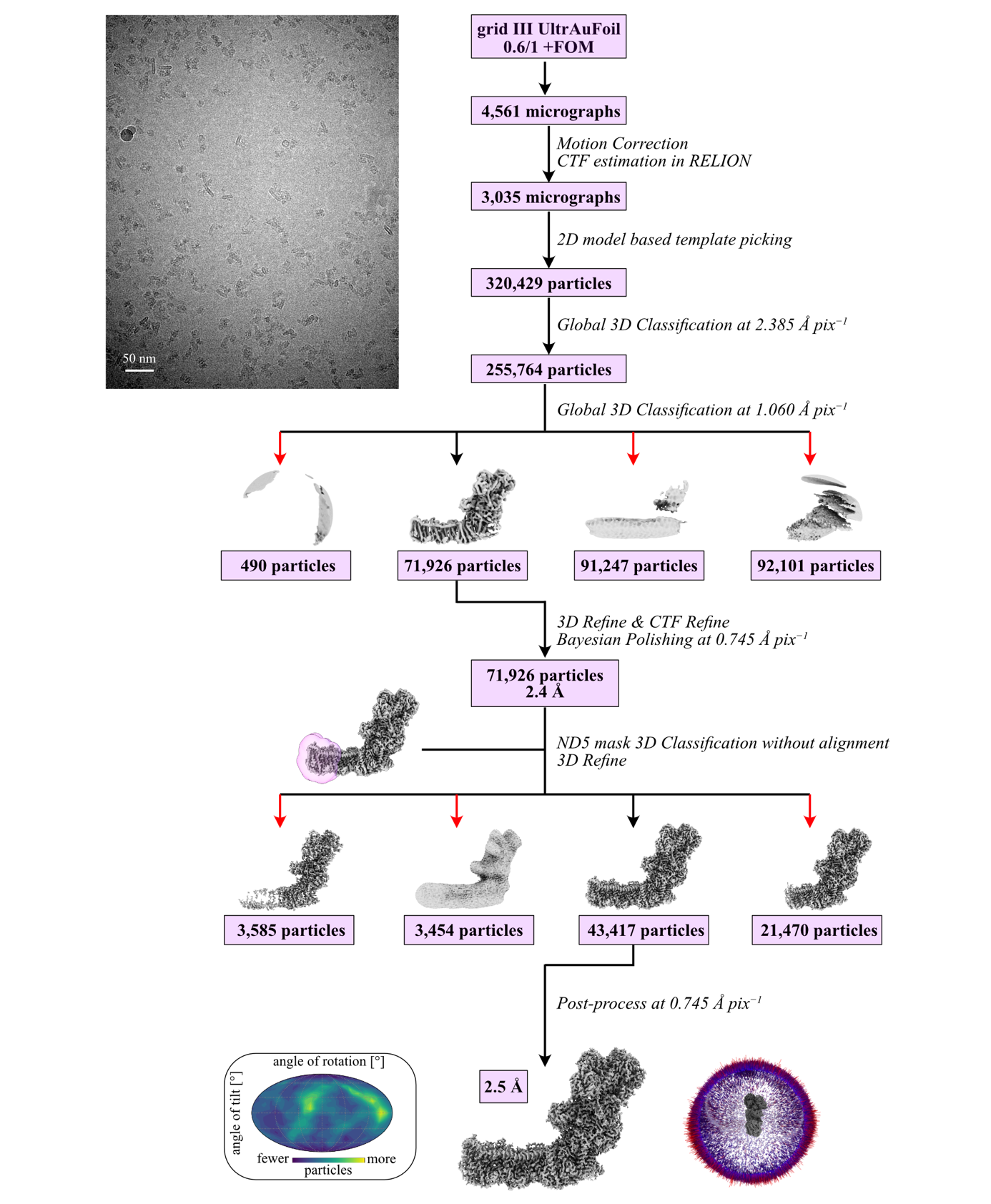

**Figure S4. Cryo-EM data processing pipeline for grid 3 for *Pd*-CI-ND in the presence of FOM.** The red arrows indicate particles that were excluded, and the orientation distributions are shown.

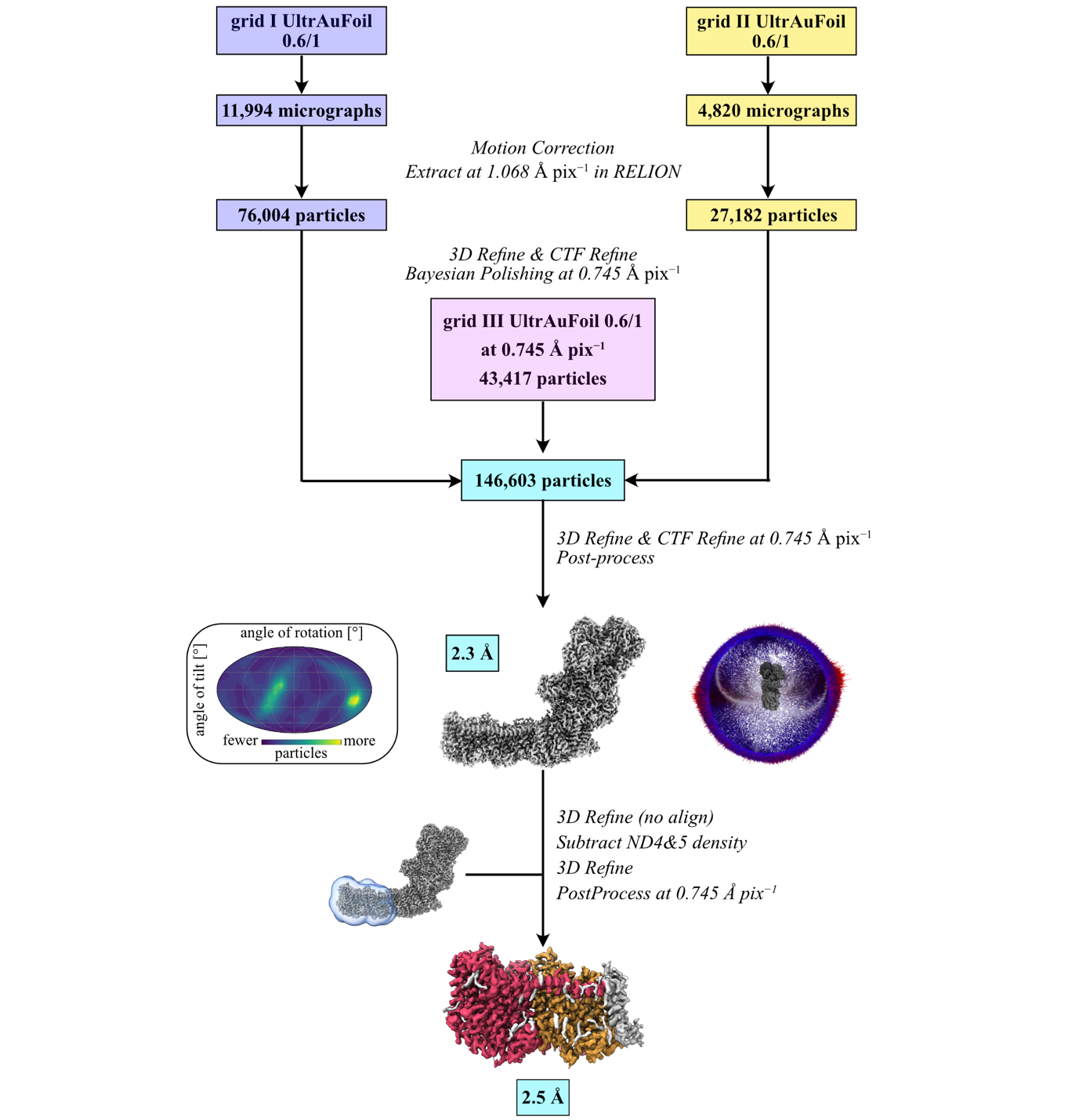

**Figure S5. The processing pipeline to combine data from cryo-EM grids 1, 2 and 3 for *Pd*-CI-ND, and the focussed refinement of the distal membrane subdomain.** The distal section contains subunits ND4 (orange) and ND5 (red).

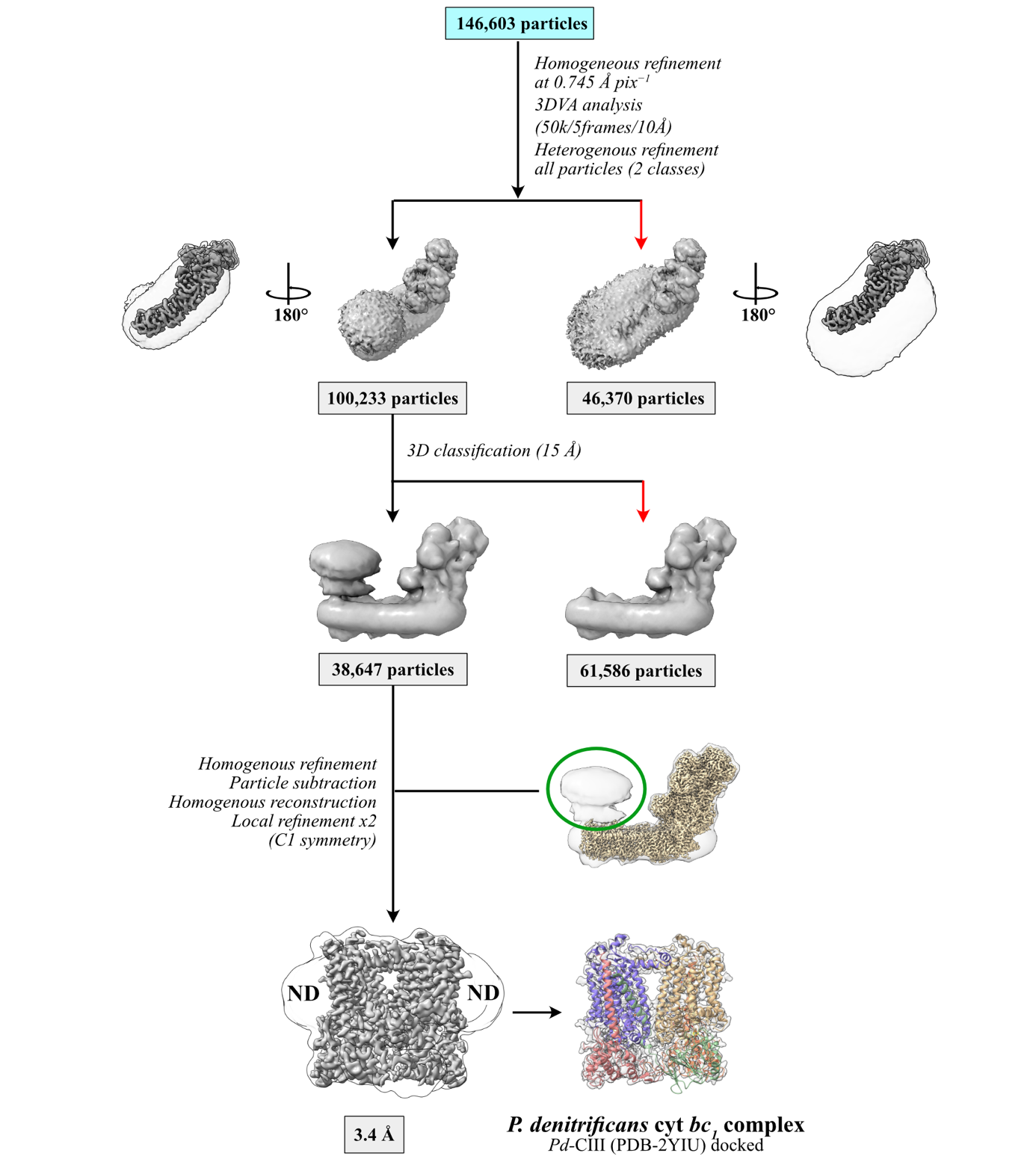

**Figure S6. Identification of two nanodisc populations and subclassification to identify the *bc_1_* complex.** Small and large NDs are indicated in solid (top-view) and transparent (bottom-view) densities. The red arrows indicate excluded classes. The structure of the *P. denitrificans* cyt *bc_1_* complex PDB-2YIU is shown fitted into the 3.4-Å density map (EMDB: 19977).
